# Multidimensional encoding of movement and contextual variables by rat globus pallidus neurons during a novel environment exposure task

**DOI:** 10.1101/2021.08.01.454624

**Authors:** Noam D. Peer, Hagar G. Yamin, Dana Cohen

**Author notes:** Corresponding author: Dana Cohen; The Leslie and Susan Gonda Multidisciplinary Brain Research Center, Bar-Ilan University, Ramat-Gan 52900, Israel.

## Abstract

The basal ganglia (BG) play a critical role in a variety of functions that are essential for animal survival. Information from different cortical areas propagates through the BG in anatomically segregated circuits along the parallel direct and indirect pathways. We examined how the globus pallidus (GP), a central nucleus within the indirect pathway, encodes input from the motor and cognitive domains. We chronically recorded and analyzed neuronal activity in the GP of rats engaged in a novel environment exposure task. GP neurons displayed multidimensional responses to movement and contextual information. A model predicting single unit activity required many task-related variables, thus confirming the multidimensionality of GP neurons. In addition, populations of GP neurons, but not single units, reliably encoded the animals’ locomotion speed and the environmental novelty. We posit that the GP independently processes information from different domains, effectively compresses it and collectively conveys it to successive nuclei.

## Introduction

The basal ganglia (BG) are a group of subcortical nuclei that are known to be involved in a variety of functions spanning the limbic, associative and sensorimotor domains. Information from different areas in the neocortex reaches the BG’s primary input structure, the striatum, where it is organized topographically and is processed by segregated circuits with minor overlap (Peters et al., 2021, Hintiryan et al., 2016, Alexander et al., 1986). From the striatum, the information is conveyed in the parallel direct and indirect pathways. Both pathways project to the output structures of the BG; however, information in the indirect pathway undergoes additional processing in the globus pallidus (GP) and the subthalamic nucleus (STN). The latter provides additional input to the BG.

It has been shown anatomically that the segregated pattern of information flow described in the striatum is maintained in the GP (Foster et al., 2020) even though the number of striatal efferents is an order of magnitude larger than that of the GP (Oorschot, 1996). By contrast, single GP neurons have been shown to respond to a variety of motor and non-motor task attributes possibly because these neurons have a very large dendritic arbor which can branch across sub-regions and therefore integrate information from several functionally distinct areas (Sadek et al., 2007, Kita and Kitai, 1994). For example, these neurons are known to encode movement direction including the signaling of movement onset, the end of a movement in a movement sequence, and are affected by static and/or dynamic load (Dodson et al., 2015, Gu et al., 2020, Brotchie et al., 1991a, Brotchie et al., 1991b, Mitchell et al., 1987, Arkadir et al., 2004). They also encode cognitive and limbic information such as the contextual setting, the relative difficulty of the task, strategies for behavioral inhibition, reversal learning and reward probability and prediction (Saga et al., 2013, Schechtman et al., 2016, Joshua et al., 2009, Gu et al., 2020, Brotchie et al., 1991b, Arkadir et al., 2004, Bischoff-Grethe et al., 2015, Lilascharoen et al., 2021). However, it remains unclear whether the GP processes all the features encoded in the striatum, or whether it extracts a few relevant features, and whether, like the striatum, these features are encoded by segregated pathways, or alternatively, are integrated by single GP neurons.

When exposed to an unfamiliar environment, animals typically initiate exploratory behavior; i.e., they engage in rapid movement throughout the environment, typically starting with its perimeters and then extending into the center. This behavior allows animals to gradually familiarize themselves with the environment. Two distinct processes are essential for successful task performance: (1) a cognitive distinction between familiar and unfamiliar environments, and (2) the planning and execution of the physical locomotion during the familiarization process. These components are dissociable because the internally generated locomotion undergoes habituation as the animal familiarizes itself with the environment, whereas the identity of the environment remains unchanged as long as the animal stays in a given space. Hence, the novel environment exposure (NEE) task is well suited for studying how the GP integrates and extracts information encompassing different domains.

Previously, we showed that non-overlapping populations of striatal projection neurons – the medium spiny neurons – reliably encode the locomotion and environment identity during performance of the NEE task (Yamin et al., 2013). Here, we inquired whether during a similar task, GP neurons would encode either the animal’s actions or the environmental identity, or both. We chronically recorded and analyzed the activity from rat GP neurons with respect to locomotion, rearing up on the hind limbs, grooming and the different environments while the animals performed the NEE task. We show that populations of GP neurons reliably encode the animals’ average locomotion and the environmental novelty. In contrast, single neurons encoded both attributes poorly even though the majority of the GP neurons (90 %) responded significantly to at least one task variable. The likelihood of a GP neuron to respond to a given task-related variable was independent of whether that neuron responded to other variables. Concomitantly, using generalized linear model, we found that many behavioral variables contributed to the prediction of single neurons’ firing rate traces during the task. These findings suggest that GP neurons integrate different types of task-relevant information, and distribute it independently across populations of single neurons.

## Materials and Methods

### Animals

All procedures were approved by the Bar Ilan University Institutional Animal Care and Use Committee and were performed in accordance with the National Institutes of Health guidelines. Data were collected from 7 Long Evans rats (3–7 months old). All animals were housed two in a cage, separated by a divider after surgery, on a 12/12 h light/dark cycle and had *ad-libitum* access to food and water. Experiments were performed during the light phase.

### Surgery

The surgical procedure has been described in detail elsewhere (Benhamou, Bronfeld et al. 2012). In brief, 7 adult male Long Evans rats weighing 460-545 gr (bred in-house) were initially sedated with 5% isoflurane and then injected intramuscularly with ketamine HCl and xylazine HCl (100 and 10 mg/kg, respectively). Supplementary injections of ketamine and xylazine were given as necessary. The skull surface was exposed and two 0.5 × 2 mm^2^ craniotomies were made above the GP. Each craniotomy was shaped as a rectangle centered on (anterior-posterior: 1.65 mm; mediolateral: ± 4.1 mm; specific coordinates were: [−0.7, ± 3.7], [−1.0, ± 3.3], [−2.6, ± 4.5], [−2.3, ± 4.9] mm relative to Bregma; (Paxinos, 2013)). Custom made microwire electrodes (microwires of S-isonel-coated tungsten, California Fine Wire Company, 50 μm in diameter arranged in a 2×8 array) were lowered from the surface of the brain while recording neural activity. The final placement of the electrodes was determined based on the measured coordinates from the surface of the brain (5.5-6.6 mm) and the quality of neural activity. Electrode placement in the GP was confirmed histologically after electrolytic lesioning, perfusion with 10% formalin, brain fixation with 20% sucrose, sectioning and acetylcholinesterase (AChE) staining. A protocol for AChE staining was obtained from neurosciencecourses.com and adjusted for perfused tissue. The rats were given at least 1 week to recover prior to recording.

### Novel Environment Exposure (NEE) task

24 h prior to the task, each rat was individually housed in a clean cage (35 x 46 x 20 cm^3^) with a small amount of bedding to familiarize the animal with its cage. Then the cage lid was opened, and the rat was connected to the head stage for at least 30 minutes before the beginning of the experiment to allow for habituation to the recording wires and the experimenter’s presence. Each recording session comprised 15 minutes of baseline activity in the home cage (BL), 15 minutes in the novel environment (NE) which was a clean cage identical to the home cage in size and amount of bedding but lacking familiar odors, and an additional 15 minutes back in the home cage (HC). The short transition periods between environments were removed from the analysis. Sessions were repeated once a week for up to 4 consecutive weeks. The animals’ behavior was monitored by a digital camera synchronized with the recording system (30 frames/s, Cineplex, Plexon Inc.).

### Data acquisition

Neuronal activity was recorded throughout the performance of the NEE task. The activity was amplified, band-pass filtered at 0.5-8000 Hz and continuously sampled at 40 KHz using the OmniPlex data acquisition system (Plexon Inc). The broadband activity was band-pass filtered for spiking activity (300-8000Hz). Offline sorting (offline sorter, Plexon Inc) was performed on all recorded units and only single neurons were taken for further analysis in Matlab (R2013b, MathWorks Inc., Natick, MA).

### Data analyses

#### Behavior analysis

The animals’ behavior was video recorded during the NEE task. First, the grooming and rearing up times were manually noted from the video. Then, for all other frames, the coordinates of the animal’s center of mass were extracted using CinePlex Studio recording & tracking software (Plexon inc), and convoluted with a 9 frame Gaussian window. The speed of the animal in each frame was calculated by measuring the distance travelled in two consecutive frames and dividing it by the inverse of the frame rate. The animal’s speed was averaged in 0.5 s bins. Bins in which the animal’s speed was above 5 cm/s were considered locomotion bins. Bins in which the animal’s speed was lower than 0.5 cm/s were considered quiet awake (QA) periods. Bins with intermediate speeds were marked as ‘others’.

#### Exploratory behavior duration

The time the animals were engaged in exploratory behavior was measured from the time they were transferred to the NE / HC until identifying 15 s without locomotion or rearing up on hind limbs.

#### Coefficient of variation (CV)

The standard deviation of the interspike interval distribution divided by its mean.

#### Action encoding neurons

For each neuron, the firing rate distribution in 0.5 s non-overlapping bins for each behavior (QA, grooming, rearing up and locomotion) was calculated for the whole session. To determine whether a neuron significantly modulated its firing rate in response to any of these behaviors, the firing rate distributions of all behaviors were compared to its firing rate distribution calculated during QA using a 1-way unbalanced ANOVA (p < 0.01).

#### Experimental condition encoding neurons

We calculated the firing rate distributions for each neuron in 0.5 s non-overlapping bins during the QA periods separately for each environmental condition (BL, NE and HC). To determine whether a neuron significantly changed its firing rate during any of the environments we compared its firing rate distributions during QA periods across environments using a 1-way unbalanced ANOVA (p < 0.01).

Neurons showing a significant ANOVA (p < 0.01) after Bonferroni correction for multiple comparisons (n=156 comparisons for behaviors and environments) were tested post-hoc to determine which action and/or environment they responded to. A neuron that responded differently during NE as compared to BL and HC was considered a novel environment encoding neuron.

#### Generalized Linear Model (GLM)

To quantify the contribution of the different variables to neural activity, we used a GLM. We used the Matlab function stepwiseglm to fit the per trial neuronal firing rate in 5 sec bins during the NEE paradigm. stepwiseglm begins with an initial constant model and takes forward or backward steps to add or remove variables, until a stopping criterion is satisfied. We used a linear model without interaction terms and the Akaike information criterion to add or remove predictors and assumed a Poisson distribution of the firing rate. The predictors included continuous and categorical variables. The continuous variables quantified kinematics including speed and acceleration (calculated as the derivative of the speed). Both predictors were z-scored prior to modeling. The categorical variables were an action vector corresponding to the per bin main activity of the animal (QA, grooming, rearing up, locomotion, or others as defined in the behavioral analysis), and an experimental condition vector corresponding to the per bin experimental condition (BL, NE or HC). The coefficient of determination, *R^2^* was used as a measure of the models’ accuracy. To evaluate the significance of each model, we calculated the *R^2^* of 25 shifted models, in which the predictors’ matrix was shifted forward in time by n bins of 5 s (n = 1, 2,…, 25 bins) relative to the firing rate which remained unchanged. The model was considered significant if its *R^2^* exceeded the 95% confidence intervals of the *R^2^* distribution of the shifted model (Harris, 2021).

#### Linear regression analysis

Linear regression was calculated between the firing rates of one individual neuron at a time or for all the neurons (in 5 s bins) and a target vector – either the speed of a single trial (for single neurons) or the average speed over all trials (for the population of neurons) or the NE condition. The target vector for the NE condition contained ones at the times the animal was at the NE and zeroes at all other times (identically for single neurons and the population). The regression was estimated using the coefficient of determination *R^2^*. To control for the large number of degrees of freedom we compared the calculated *R^2^* value to that of a bootstrap consisting of 100 different shuffles of each neuron’s firing rate.

#### Cross Correlation

Cross-correlograms (CC) were calculated for ±1 s in 1 ms bins for all neuronal pairs that were not recorded from the same electrode (Bar-Gad et al., 2001). A total of 147 CCs from 15 sessions were calculated and analyzed for significance. A CC was considered significant if 3 consecutive bins in the inner part of the CC (±100 ms) crossed the 95 % confidence intervals calculated from the outer parts of the CC (> |±500ms|) (Nevet et al., 2007, Oliveira-Maia et al., 2012, Kopelowitz et al., 2014)

### Statistical analysis

All the data are presented as mean ± SEM unless otherwise noted. All statistical tests were ANOVAs unless otherwise noted. All post-hoc corrections for multiple comparisons were Bonferroni corrections.

## Results

### Rats explore the novel environment by locomotion and rearing up on their hind limbs

Seven male Long-Evans rats engaged in the NEE task (see Methods) once a week for up to four consecutive weeks. Each experimental session comprised 15 minutes in the home cage, defined as baseline activity (BL), after which the animal was transferred to the novel environment (NE), a cage identical to the home cage but lacking familiar odors or objects. After a period of 15 minutes in the novel environment, the animal was transferred back to its home cage (HC) for an additional 15 minute period (Fig. 1A). Upon transition to the NE, the rats initiated exploratory movement that consisted primarily of locomotion and rearing up on their hind limbs and leaning against the cage wall (see example session in Fig. 1B). The exploration of the NE gradually ceased and the animals tended to select a preferred corner and remained motionless while sporadically initiating short epochs of exploratory behavior. When transferred back to their home cages the animals reinitiated exploratory behavior; however, the exploration time of the HC was significantly shorter than for the NE (326 ± 53 s and 475 ± 66 s for HC and NE, respectively, paired t-test, t = 2.37, df = 26, p = 0.026, see Methods). Along with the increased exploration time, the fraction of time the animals spent walking and rearing up was significantly higher in the NE as compared to the BL and the HC (Fig. 1C, locomotion: 5.5 ± 0.8 %, 9.9 ± 1.0 % and 7.1 ± 0.8 % for BL, NE and HC, respectively; ANOVA, F = 7.06, df = 2, p = 0.0015; Fig. 1D, rearing up: 3.9 ± 0.8 %, 16.2 ± 2.4 % and 7.1 ± 1.6 % for BL, NE and HC, respectively; ANOVA, F = 13.08, df = 2, p = 1.8 x10^-5^, post-hoc testing with multiple comparison adjustments). These findings suggest that rats use a strategy of interleaved locomotion and rearing up on their hind limbs to explore the NE.

**Figure 1.**
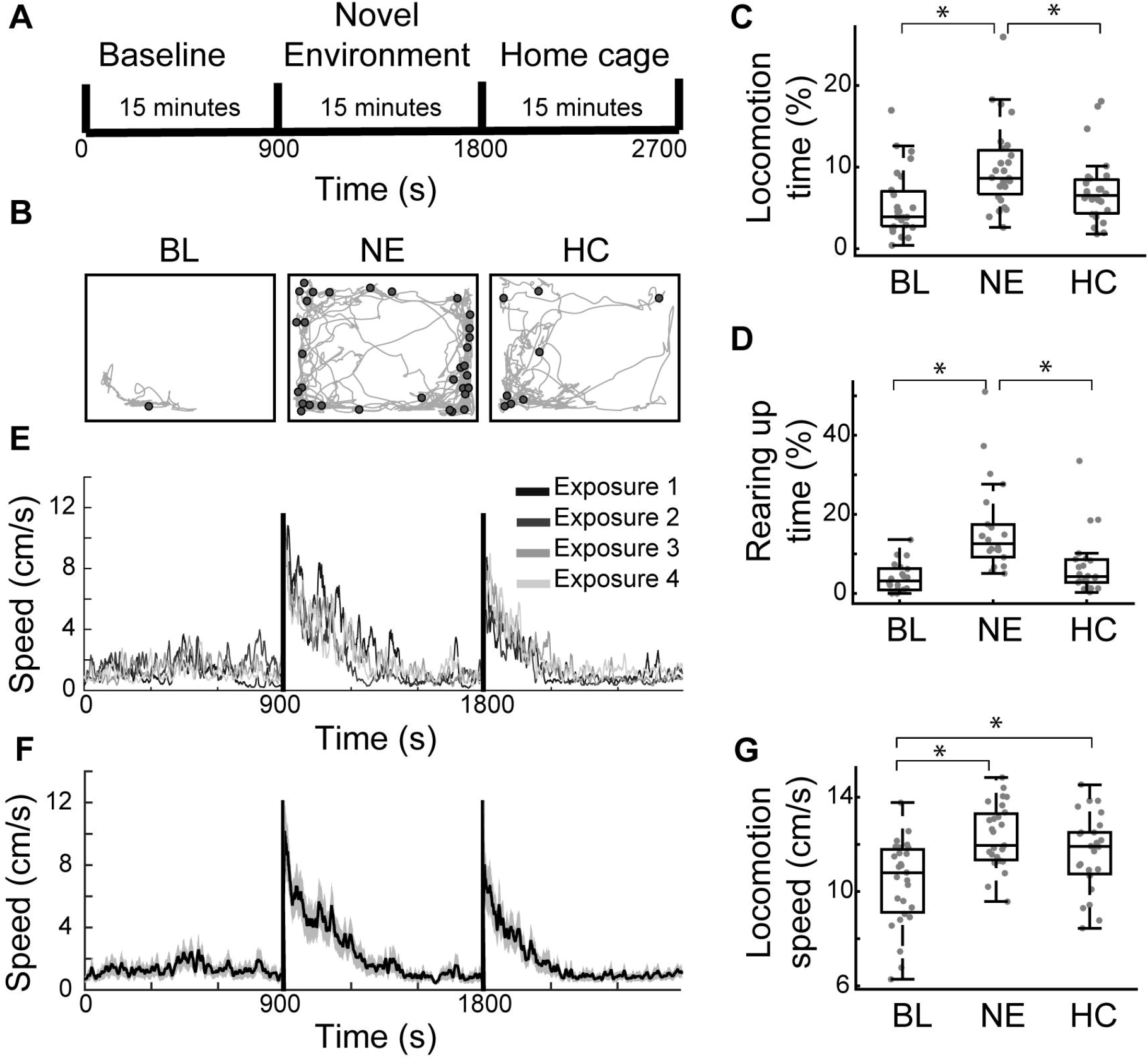
Rats explore the novel environment by locomotion and rearing up on their hind limbs. **A.** Experimental timeline: rats spent 15 minutes in their home cage, defined as the baseline activity, then were transferred to a novel environment for 15 minutes, and finally transferred back to their home cage for another 15 minutes. **B.** A single animal’s trajectory (grey lines) and rearing up locations (circles) during BL (left), NE (center) and HC (right) of an example session. Rectangles represent the cage (35 x 46 cm^2^). **C.** Boxplots of the fraction of time (in %) the animals spent in locomotion in each environment. In each box, the central mark indicates the median, and the bottom and top edges of the box indicate the 25th and 75th percentiles, respectively. The whiskers extend ± 2.7 SD. Grey dots denote individual sessions. **D.** Boxplots of the fraction of time (in %) the animals spent rearing up in each environment. Box plot conventions as in (C). **E.** Animals’ speed in 5 s bins averaged across sessions from the same exposure week (1 - 4). Grey scale denotes the exposure number. Black vertical lines represent transition times between experimental conditions. **F.** Animals’ speed in 5 s bins averaged across all sessions (black) and SEM (grey shading). Black vertical lines represent transition times between experimental conditions. **G.** Boxplots of the animals’ speed for each experimental condition. Box plot conventions as in (C). In all subplots, the star denotes an ANOVA, p < 0.01.

As reported in mice (Yamin et al., 2013), the rats did not habituate to the task and displayed a similar pattern of locomotion speed in all sessions irrespective of the number of exposures (Fig. 1E). Hence, locomotion speed was averaged across sessions (Fig. 1F). Movement speed was significantly higher in the NE and the HC as compared to BL (Fig. 1G; 10.4 ± 0.3 cm/s, 12.3 ± 0.2 cm/s and 11.6 ± 0.3 cm/s for BL, NE and HC, respectively; ANOVA, F = 10.56, df = 2, p = 9 x10^-5^, post-hoc testing corrected for multiple comparisons). The pattern of the average locomotion speed was similar to that previously reported in mice, suggesting that epochs of rearing up were cancelled out.

### Single GP neurons display multidimensional responses during the NEE task

We chronically recorded the activity of 78 neurons in the GP of 7 rats during performance of the NEE task (n = 27 sessions). The electrodes’ position in the GP was verified histologically for all recorded neurons (see Methods). During BL, GP neurons displayed an average firing rate of 5.37 ± 0.49 spikes/s and a CV of 0.96 ± 0.05. Throughout the task, the neurons displayed tonic firing with complex modulations (see two example neurons in Fig. 2A and 2B; see movie). Examination of these firing rate modulations with respect to different animal behaviors (locomotion, rearing up and grooming, denoted in the figure by different shades of green) indicated that the neuron shown in Fig. 2A increased its firing rate for locomotion and rearing up and decreased its firing rate during grooming as compared to its firing rate during the QA periods (Fig. 2C; see Methods). The neuron shown in Fig. 2B increased its firing rate during grooming and rearing up but not during locomotion as compared to the QA periods (Fig. 2D). Examination of firing rates measured in these example neurons during the QA periods in the different environments showed that the neuron in Fig. 2A displayed a higher firing rate in the NE and the HC as compared to BL, whereas the neuron shown in Fig. 2B did not distinguish between environments (Fig. 2E and 2F, respectively).

**Figure 2.**
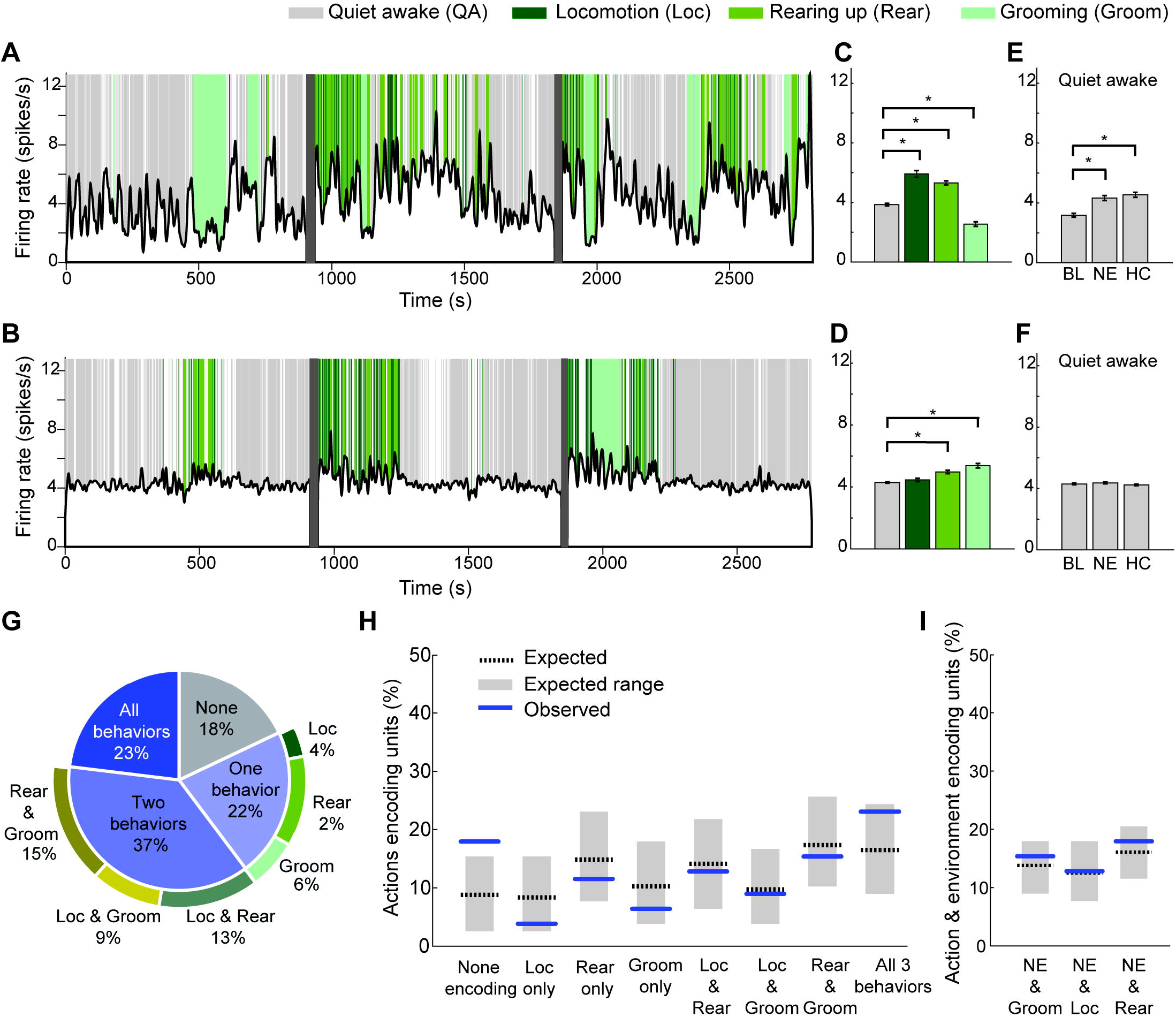
Single GP neurons display multidimensional responses during the NEE task. **A.** An example firing rate trace of a neuron (black) throughout the NEE task, calculated in 0.5 s bins and smoothed using a Gaussian window of 40 bins. Background colors represent the activity of the animal and are denoted in the figure legend. Grey vertical lines represent transition times between experimental conditions. See also movie. **B.** Same as in (A) for a different neuron. **C.** The average firing rate of the neuron plotted in (A) while the animal was engaged in the different actions. Error bars are SEMs. Color scheme as in (A). **D.** Same as (C) for the neuron plotted in (B). **E.** The average firing rate during quiet awake periods for the different experimental conditions. Same neuron as in (A). Error bars are SEMs. **F.** Same as (E) for the neuron plotted in (B). **G.** Pie chart of the proportion of neurons encoding different combinations of tested actions. **H.** The expected (dotted black line) and the observed (blue line) percentages of units encoding different possible combinations of actions. The expected value for these combinations was calculated from the marginal probabilities of the tested actions assuming independent processes. Grey rectangles represent two-way 99 % confidence intervals calculated from 10,000 repetitions of random draws of responses from the marginal probabilities of each action, and were plotted for demonstration sake only. **I.** The expected (dotted black line) and the observed (blue line) percentages of units encoding both the environmental novelty and any of the analyzed actions. The expected values were calculated from the marginal probabilities to encode an action and the NE assuming independent processes. Grey rectangles as in (H) but for the NE and the different actions. In all subplots the star denotes ANOVA, p < 0.01.

Quantification of firing rate alterations in the whole population during the different motor behaviors showed that the majority of GP neurons displayed significant firing rate alterations in response either to one (21.8 %), two (37.2 %) or three behaviors (23.1 %) whereas only a minority (17.9 %; Fig. 2G) of the neurons did not respond to any of these motor behaviors. To examine whether the likelihood of a GP neuron to encode one motor behavior co-varied with its likelihood to encode another behavior, we calculated the probability of GP neurons to encode different combinations of behaviors based on their marginal probabilities to encode single behaviors. The use of the marginal probabilities sufficed to predict the probabilities of GP neurons to encode a single behavior, two behaviors or three behaviors (Fig. 2H; *χ*^2^ test for independence, *χ*^2^ = 3.4, df = 4, p = 0.49). This indicates that the likelihood of a GP neuron to encode a given motor behavior was independent of whether or not it encoded other behaviors. That is, even though locomotion and rearing up are part of the exploration process and rearing up and grooming have a common body posture, their joint probability was similar to that expected from independent processes.

Quantification of firing rate alterations of GP neurons during the QA periods in the different environments (experimental conditions; see Methods) showed that the majority (55%) of the neurons encoded information about the environment, out of which 46.5% (20/43) distinguished the NE from the other two environments. Only 8/78 (10.3 %) of the neurons did not respond to any task attribute. We then examined whether the likelihood of a GP neuron to encode any of the tested motor behaviors co-varied with its likelihood to encode the NE. To that end, we calculated the likelihood of GP neurons to encode the NE and/or any motor behavior by using the marginal probabilities of the different behaviors and the NE. The marginal probabilities sufficed to predict the probability of the GP neurons to encode any behavior and/or the NE (Fig. 2I; *χ*^2^ test for independence, *χ*^2^ = 0.15, df = 1, p = 0.70, *χ*^2^ = 0.23, df = 1, p = 0.63, *χ*^2^ = 0.87, df = 1, p = 0.35, for grooming, locomotion and rearing up, respectively). Thus, single GP neurons appear to be able to encode multiple task-related variables, and the propensity to encode one variable is independent of the neuron’s propensity to encode other variables.

### Many behavioral predictors contribute to the activity of single GP neurons

To directly assess the contribution of the different behavioral variables to the activity of single GP neurons, we examined whether the firing rate profile of each neuron could be predicted by a generalized linear model (GLM) which took into account a variety of task-related variables (Engelhard et al., 2019). The model included continuous and categorical variables (see Methods). The continuous variables were the speed and the acceleration of the animal’s center of mass. The categorical variables consisted of the type of behavior and the experimental conditions. The behavioral variables included bins tagged according to whether the animal was engaged in grooming, rearing up, locomotion or QA. The remaining bins were marked as ‘others’. The experimental condition variables were composed of bins tagged according to the environment in which the animal was located (i.e. BL, NE or HC). The variables were fed into the GLM to predict the firing of single GP neurons recorded during the NEE task (Fig. 3A). We used *R*^2^ as a metric to assess the model’s accuracy. The significance of each model was evaluated by testing whether its accuracy exceeded the confidence interval of an accuracy distribution calculated from control models in which the neuron’s firing was predicted from the same behavioral data that were shifted forward in time (see Methods). The GLM of 40/78 (51.3 %) of GP neurons were significant. Two examples of the estimated firing traces (blue) overlaid on the actual firing traces of their corresponding GP neurons (black) are shown in Fig. 3B and 3C. Fig. 3D depicts the distributions of the R^2^ values of the significant (blue; range: 0.03 - 0.5) and non-significant (grey; range: 0.001 - 0.34) models.

**Figure 3.**
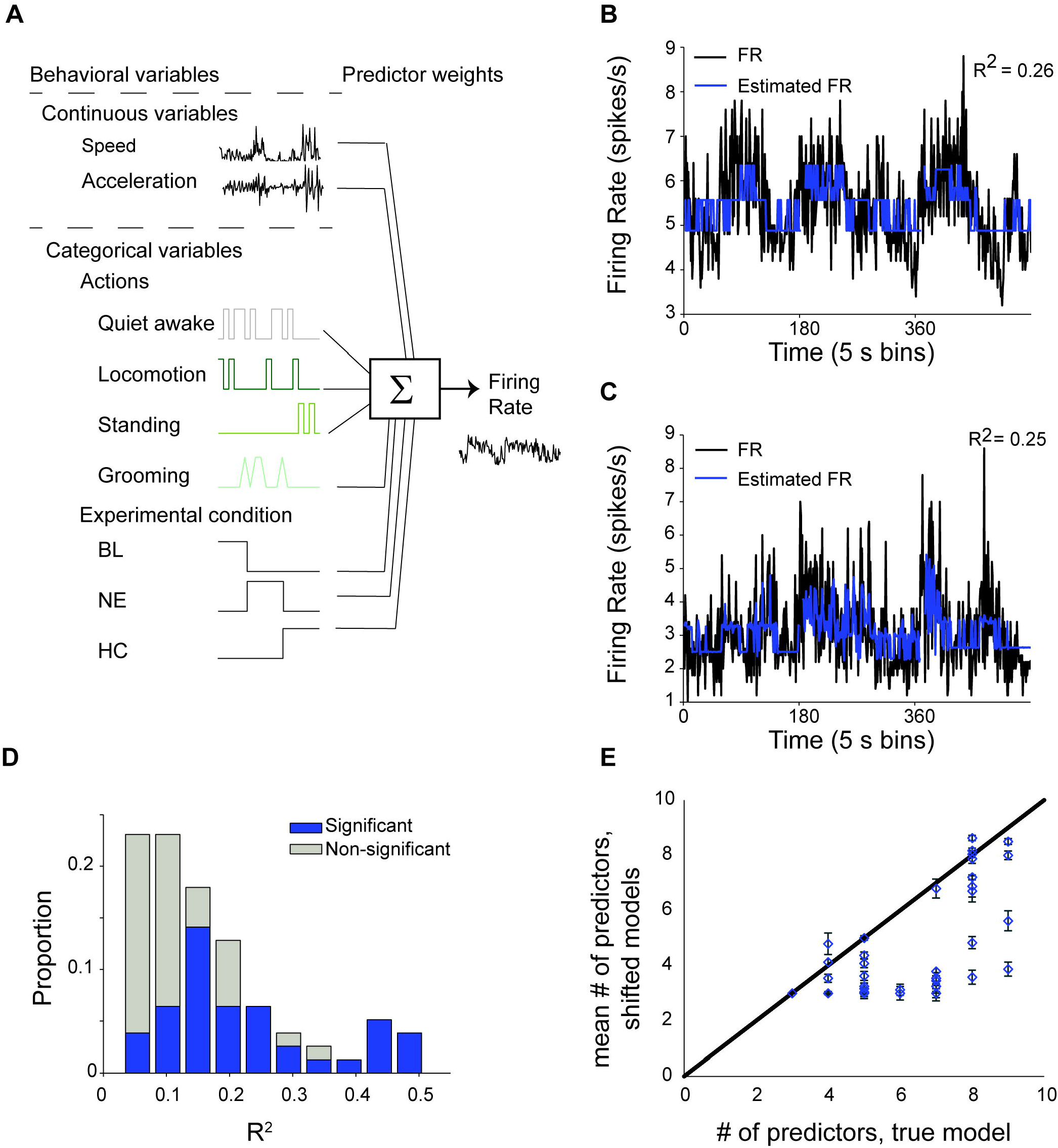
Many behavioral predictors contribute to the activity of single GP neurons. **A.** Schematic illustration of the behavioral and environmental variables used by the GLM to predict single neuron’s firing rate. **B.** An example of a GLM fit (blue) to the actual firing rate trace of a single neuron (black). **C.** Same as (B) for a different neuron. **D.** The distribution of *R*^2^ values for all GLMs. Blue and grey denote the significant and the non-significant models, respectively. **E.** The number of predictors in the true GLM (x axis) versus the mean ± SEM number of predictors in a set of shifted GLMs (y axis; see Methods). Only significant models are plotted.

The number of contributing predictors to the significant GLMs was high (6.32 ± 0.28) considering that each model could potentially have a maximum of 9 variables. In the vast majority (33/40) of the significant models, the number of contributing predictors of the neurons’ firing rates was higher than that of the shifted models (Fig. 3E; 4.81 ± 0.07 predictors in the shifted models, KW test, H = 24.62, df = 1, p = 6.9 x 10^-7^). Thus, many task-relevant variables appeared to contribute to the firing of each GP neuron. This outcome is consistent with the result described in the previous section that GP neurons modulate their firing rate significantly in response to behavioral and environmental conditions.

### The animals’ locomotion and the environmental novelty are encoded by populations of GP neurons but not by single neurons

The fact that many task-relevant variables were required for successful modeling of single neuron activity implies that input about these variables may be distributed across the population rather than be encoded by single neurons. To investigate this possibility, we used a linear regression model to approximate two task attributes that were common to all animals: the average locomotion speed and the environmental novelty (see Methods). The population of GP neurons yielded high *R*^2^ values for the animals’ average speed (*R*^2^ = 0.77, Fig. 4A) and the environmental novelty (*R*^2^ = 0.70, Fig. 4B). To control for the degrees of freedom, we randomly shuffled the firing rate of the neurons 100 times and recalculated the models’ accuracy (see Methods). The probability to achieve the population’s *R*^2^ values or higher using the shuffled data was 0 for both the locomotion speed and the environmental novelty (Fig. 4C & 4D, respectively) suggesting that the high performance of the regression model was not due to the number of predictors (n = 78 neurons).

**Figure 4.**
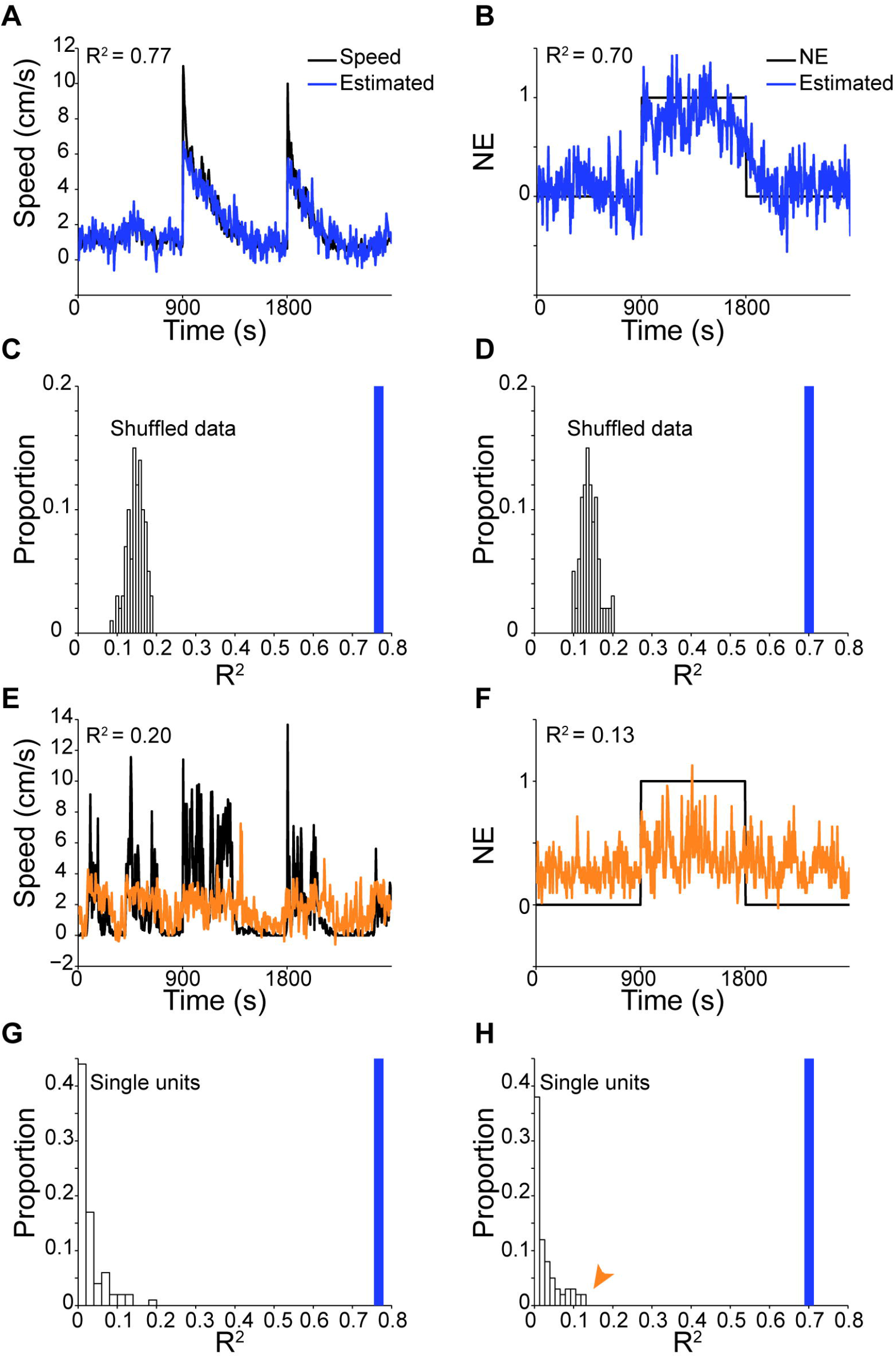
The animals’ locomotion and the environmental novelty are encoded by populations of GP neurons but not by single GP neurons. **A.** The animals’ speed in 5 s bins averaged across all sessions (black), and the estimated speed (blue) calculated by a linear regression model using the firing rate of all recorded neurons (n = 78). **B.** Same as (A) for NE identity. **C.** The distribution of *R*^2^ values of linear regression models applied to shuffled firing rates to predict the animals’ average speed (see Methods). Blue line is the R^2^ value of the true model. **D.** Same as (C) for estimating the NE identity. **E.** The best example of a linear regression model applied to a single GP neuron to estimate the animal’s speed. The estimated speed (orange) overlaid on the animal’s speed in 5 s bins (black). **F.** Same as (E) for estimating the NE identity. **G.** The distribution of R^2^ values of linear regression models applied to single units to predict the animal’s speed in a session. Blue line is the R^2^ value of the model using all neurons (n = 78). The example shown in E is marked by an orange arrowhead. **H.** Same as (G) for estimating the NE identity. The example shown in F is marked by an orange arrowhead.

The high performance of the regression model on the population could have emerged either from a few single neurons that effectively predicted the two task attributes, or alternatively, that the minor contribution of many single neurons were summed. To distinguish between these possibilities, we applied linear regression models to single GP neurons to assess how well they encoded the animal’s speed and presence in the novel environment. Unlike the population, single neurons approximated both task attributes poorly (the best example models are shown in Fig. 4E and 4F, for speed and environmental novelty, respectively). Fig. 4G & 4H depict the *R*^2^ distribution of single units (white bars) as compared to the population (blue bar), suggesting that each neuron by itself made a minor contribution to the population model. The gain in regression performance when taking into account the population as compared to single units could have emerged from averaging across correlated signals with random noise, thus enhancing the signal to noise ratio. To control for this possibility, we calculated the cross-correlograms (see Methods) of all simultaneously recorded pairs of GP neurons. We found that a minority of the tested pairs (12/147) were significantly correlated. This suggests that individual neurons may convey nonoverlapping idiosyncratic information.

## Discussion

This study explored the encoding of motor and cognitive attributes of the NEE paradigm by GP neurons. We showed that rats explored the novel environment by locomotion and rearing up on their hind limbs. Single GP neurons encoded behaviors performed by the rats such as grooming, rearing up and locomotion and also distinguished between environmental conditions including identifying the novel environment. The probability of a GP neuron to encode one or more of these task attributes was independent of whether the neuron encoded other attributes, and the activity of simultaneously recorded pairs of neurons was uncorrelated. Prediction of the single neuron firing rate by GLM demonstrated that many behavioral variables contributed to each model. Nevertheless, single GP neurons failed to approximate either the animal’s locomotion or the animal’s presence in the novel environment. By contrast, both task attributes were reliably encoded by a population of GP neurons. These data indicate that task-relevant information is distributed across GP neurons in a non-overlapping manner, and can therefore be extracted from the population, rather than from single GP neurons.

It has been shown that single neurons in the globus pallidus external segment (GPe) of primates, which is homologous to the rodent GP, can encode more than one task-relevant variable. For example, GPe neurons were reported to encode the direction of reaching movement and reward during a probabilistic visuomotor task (Arkadir et al., 2004), and could encode effort and reward in a task where a different force was required to obtain different amounts of reward (Nougaret and Ravel, 2018). These task-relevant parameters were integrated linearly, suggesting their independent encoding by GP neurons (Arkadir et al., 2004). Moreover, GP neurons were reported to generate dynamic ensembles which at any given time encoded the most relevant information for best task performance (Nougaret and Ravel, 2018, Adler et al., 2012, Saga et al., 2013). Our results in the rodent GP are in agreement with those described in the primate GPe; and extend them by showing that single GP neurons encode different movements as well as the nature of the environment simultaneously and independently, thus encompassing both the motor and cognitive domains.

During the NEE task, striatal MSNs encoded either the animal’s locomotion or the environmental novelty but not both (Yamin et al., 2013). The identity of MSN encoding each task attribute was not addressed. Hence, it remains unclear whether these task attributes were encoded selectively by MSNs from either the direct or the indirect pathway, or both. The fact that GP neurons responded to both task attributes indicates that MSNs from the indirect pathway contributed to task performance. Furthermore, it rules out the possibility that the GP processed these task attributes in segregated parallel pathways. It also remains unclear whether MSNs from the direct pathway also process the two task attributes, and whether the GP selects specific task-relevant features to maximize performance or optimally compresses all the converging data from the striatum using dimensionality reduction (Bar-Gad et al., 2000, Bar-Gad et al., 2003b). The NEE paradigm may be too simplistic to differentiate between these two alternatives because both the motor and the cognitive features are essential for task performance. Hence, deciphering whether GP neurons are tuned to extract functionally relevant features, or to compress all converging data from the striatum may require tasks utilizing more complex feature representations.

A recent study of BG networks showed that the mouse GP faithfully follows the striatal topography and is characterized by a small percentage of converging inputs arising from different regions of the striatum (Foster et al., 2020). This regional specificity, which is more segregated than the coarse division into limbic, associative and sensorimotor domains, highlights the expectation of a limited overlap in converging afferents onto the GP. Hence, it remains unclear how GP neurons display multidimensional, non-overlapping and uncorrelated encoding of task attributes in both primates and rodents. GP neurons have a large dendritic arbor extending across large portions of the GP volume that is aligned perpendicularly to MSN axons (Kita and Kitai, 1994, Park et al., 1982, Jaeger and Kita, 2011, Percheron et al., 1984). This anatomical organization may allow GP neurons to integrate information from relatively distant striatal domains but at the same time yield a substantial overlap of inputs in neighboring GP neurons. The high overlap in converging inputs is likely to generate interactions between the probabilities of GP neurons to encode different task attributes and enhance neuronal cross-correlations as shown here, and has been reported to be weak to nonexistent (Bar-Gad et al., 2003a). Two distinct scenarios may account for these phenomena: the first utilizes the high convergence ratio of 40:1 between MSNs to GP neurons (Oorschot, 1996). In this case, MSN projections onto GP neurons are highly selective so that despite the high spatial overlap, each GP neuron receives afferents from a distinct group of MSNs. The second scenario suggests that although overlap in GP afferents exists, their common input is cancelled out by other mechanisms such as the recurrent collateral axons of GP neurons (Kita et al., 2004) which have a complex spatial spread in the GP (Matsumura et al., 1995). Another possibility is the strong reciprocal connections between the GP and the subthalamic nucleus (STN) (Atherton et al., 2013, Baufreton et al., 2009, Magill et al., 2000).

The encoding scheme utilized by GP neurons is substantially different from the one utilized by the striatal projection neurons, the MSNs: GP neurons display multidimensional responses whereas the MSNs respond typically to a single event by a short bursting activity (Fobbs et al., 2020, Berke et al., 2009, Shidara et al., 1998, Atallah et al., 2014, Jog et al., 1999, Jin and Costa, 2010). Further research is needed to determine why the GP converts the temporally precise, highly specific information present in single MSNs into spatially distributed information where each component carries little information about the task. Additional experiments are required to determine how information from the striatum and the GP is integrated in the basal ganglia output structures, the entopeduncular nucleus (EP) and the substantia nigra pars reticulata (SNr), and which task attributes are represented in these structures.

## Supporting information

Movie: Activity of an example neuron during different rat behaviors

## Acknowledgments

This research was supported by the ISRAEL SCIENCE FOUNDATION (grant No. 1786/16) and by the SYNCH project funded by the European Commission under the H2020 FET Proactive programme (Grant agreement ID: 824162).

## Author contributions

NP and DC designed the research, NP performed the research, NP, HGY and DC analyzed the data, NP, HGY and DC wrote the paper

## Declaration of interests

The authors report no conflict of interest.

## Multimedia

**Movie**: Activity of an example neuron during different rat behaviors. Same neuron as in Fig. 2A.

Related to Fig. 2

## Notes

### Competing Interest Statement

The authors have declared no competing interest.

